# TRIM72 restricts lyssavirus infection by inducing K48-linked ubiquitination and proteasome degradation of the matrix protein

**DOI:** 10.1101/2023.09.29.560081

**Authors:** Baokun Sui, Jiaxin Zheng, Zhenfang Fu, Ling Zhao, Ming Zhou

**Affiliations:** State Key Laboratory of Agricultural Microbiology, Huazhong Agricultural University, Wuhan, China; Hubei Hongshan Laboratory, Wuhan, China; Frontiers Science Center for Animal Breeding and Sustainable Production, Wuhan China; Key Laboratory of Preventive Veterinary Medicine of Hubei Province, College of Veterinary Medicine, Huazhong Agricultural University, Wuhan, China

**Keywords:** TRIM72, Lyssavirus, Matrix protein, Ubiquitination, Proteasome

## Abstract

The tripartite motif (TRIM) protein family is the largest subfamily of E3 ubiquitin ligases, playing a crucial role in the antiviral process. In this study, we found that TRIM72, a member of the TRIM protein family, was increased in neuronal cells and mouse brains following rabies lyssavirus (RABV) infection. Over-expression of TRIM72 significantly reduced the viral titer of RABV in neuronal cells and mitigated the pathogenicity of RABV in mice. Furthermore, we found that TRIM72 over-expression effectively prevents the release of RABV. In terms of the mechanism, TRIM72 promotes the K48-linked ubiquitination of RABV Matrix protein (M), leading to the degradation of M through the proteasome pathway. TRIM72 directly interacts with M and the interaction sites were identified and confirmed through TRIM72-M interaction model construction and mutation analysis. Further investigation revealed that the degradation of M induced by TRIM72 was attributed to TRIM72’s promotion of ubiquitination at site K195 in M. Importantly, the K195 site was found to be partially conserved among lyssavirus’s M proteins, and TRIM72 over-expression induced the degradation of these lyssavirus M proteins. In summary, our study has uncovered a TRIM family protein, TRIM72, that can restrict lyssavirus replication by degrading M, and we have identified a novel ubiquitination site (K195) in lyssavirus M.

**Author Summary:** Rabies, caused by lyssaviruses, most often by rabies virus, the most lethal zoonotic disease, is responsible for approximately 59000 human deaths globally each year. Lyssavirus M protein displays an essential role in lyssavirus assembly and budding. Here, we found that TRIM family protein TRIM72 directly interacts with lyssavirus M thereby promoting the K48-linked ubiquitination of M, leading to the degradation of M through the proteasome pathway, restricting lyssavirus release. Moreover, we identified a novel important ubiquitination site (K195) in lyssavirus M. Prior to our study, there had been no reports of a direct interaction between TRIM72 and viral proteins, an antiviral function of TRIM72 was proved in our study. It is possible that TRIM72 interacts and ubiquitinates other viral proteins from various viruses, which warrants further investigation. This research contributes to our understanding of how TRIM72 and other TRIM proteins play a role in defending against viral invasions and may inspire further research in this field.

## INTRODUCTION

Lyssavirus, a genus of the family *Rhabdoviridae*, is a single-stranded negative-sense RNA virus that encodes five viral proteins: nucleoprotein (N), phosphoprotein (P), matrix protein (M), glycoprotein (G) and large RNA polymerase protein (L). The lyssavirus genus is currently composed of 17 viral species [1], typical lyssavirus species include *Rabies lyssavirus* (RABV), *Australian bat lyssavirus* (ABLV), *Duvenhage lyssavirus* (DUVV), *European bat lyssavirus* (EBLV)*, Lagos bat lyssavirus* (LBV) and Mokola lyssavirus (MOKV), etc [2, 3]. Among those lyssaviruses, RABV is the most well-known lyssavirus and is responsible for causing rabies, resulting in approximately 59000 human deaths globally each year [4–6]. The RABV M plays a pivotal role in various aspects of the virus’s life cycle, including virus assembly/budding [7], and regulation of the balance between virus transcription and replication [8]. Moreover, RABV M also plays an important role in inhibiting the NK-κB pathway activation [9].

Post-translational modifications (PTMs) of proteins, which involve to the addition or removal of covalent functional groups, play a crucial role in regulating protein structure, localization and activity [10]. Common PTMs include phosphorylation, acetylation, ubiquitination, methylation, glycosylation, etc [11]. Ubiquitination is an essential PTM process that covalently conjugates single or multiple ubiquitin molecules to one or more lysine residues of a substrate protein [12]. This process is catalyzed by three types of enzymes: ubiquitin-activating enzymes (E1), ubiquitin-conjugating enzymes (E2) and ubiquitin-protein ligases (E3) [13]. The ubiquitin-proteasome system plays a crucial role in degrading abnormal proteins and short-lived proteins within cells to maintain normal biochemical processes; it consists of ubiquitin, three types of ubiquitin-associated enzymes (E1, E2 and E3) and the proteasome [14].

More than 600 E3 ligases have been characterized and are classified into 3 different classes: homologous to E6-AP carboxyl terminus (HECT), really interesting new gene (RING) and RING-between RING (RBR) [15]. RING proteins are the largest class of E3 ligases and among them the tripartite motif (TRIM) family of proteins was the largest subfamily of RINGs. TRIM proteins are involved in many biological processes, including transcriptional regulation, cell proliferation and differentiation, apoptosis, DNA damage repair, intracellular signal transduction and immune response, etc [16, 17]. TRIM proteins are characterized by the presence of an N-terminal tripartite or RBBC motif comprised of a RING domain, either one or two B-boxes (B1 and B2) and a coiled-coil (CC) domain, followed by a highly variable C-terminal domain [18, 19]. The C-terminal variable functional region of the TRIM protein plays a crucial role in substrate recognition thereby conferring functional specificity, the PRY-SPRY domain is the most common C-terminal domain among known TRIM proteins [20, 21].

As a member of the TRIM protein family, TRIM72 (also known as MG53) is a RING-mediated E3 ubiquitin ligase that consists of five domains: the RING domain, B-box domain, coiled-coil domain, PRY domain and SPRY domain [22]. The RING domain contains the E3 ubiquitin ligase activity responsible for mediating the ubiquitination of numerous proteins. For instance, TRIM72 directly interacts with Ras-related C3 botulinum toxin substrate 1 (RAC1) through its coiled-coil domain and suppresses RAC1 activity by catalyzing the Lys48 (K48)-linked polyubiquitination of RAC1 at Lys5 residue in HCC cells [23]; Additionally, TRIM72 induces ubiquitination of insulin receptor substrate 1 (IRS-1) with the assistant of an E2-conjugating enzyme UBE2H [24, 25]. The B-box domain, a zinc ion binding domain, plays a critical role in TRIM activity. A previous study has demonstrated that the B-box domain also regulates the interaction between TRIM72 and FAK8 [24]. The conserved coiled-coil domain of TRIM72 is critical for TRIM72 homodimerization [26]. Furthermore, the PRY-SPRY domain was shown to bind with Orai1 and regulate extracellular Ca^2+^ entry [27]. Consequently, TRIM72 is a multifunctional protein involved in various biochemical processes within cells. However, the relationship between TRIM72 and the viruses remains unclear.

In this present study, we present the major discovery that RABV infection induces the up-regulation of TRIM72 in neuronal cells and mouse brains. Over-expression of TRIM72 restricts the release of RABV in neuronal cells and reduces the pathogenicity of RABV in mice. We have determined that TRIM72 induces the degradation of RABV-M through the proteasome pathway. Our functional domain truncation analysis of TRIM72 indicates that the TRIM72-SPRY domain interacts with RABV-M. Furthermore, through the construction of a TRIM72-SPRY-M interaction model and mutation analysis, we have confirmed that TRIM72 ubiquitinates a specific site, K195, in M, thereby leading to its degradation. Significantly, we found that the K195 ubiquitination site is partially conserved among lyssavirus, and TRIM72 interacts with lyssavirus M proteins, ultimately resulting in the degradation of lyssavirus M proteins. Consequently, our study reveals a previously unrecognized TRIM72-ubiquitination-proteosome-mediated antiviral response and identified a novel ubiquitination site (K195) in lyssavirus M proteins.

## RESULTS

### RABV infection induces TRIM72 up-regulation in mouse brains and neuronal cells

To determine the relationship between RABV and TRIM72, we assessed the TRIM72 level in RABV-infected mouse brains and neuronal cells. For the in vivo experiments, 6-week-old C57 BL/6 mice were infected with 200 fluorescence focus units (FFU) RABV (CVS-B2c strain) via intracerebral injection. As a negative control, DMEM was injected into mouse brains. Brains were collected on the 6th day post-infection. The mRNA level and protein level of TRIM72 were analyzed by qPCR and western blotting. The mRNA level and protein level of TRIM72 were dramatically increased in RABV-infected mouse brains compared to the control group (Fig. 1A-1B). For the in vitro experiments, primary neuron cells were isolated and cultured, and then the neuron cells were infected with RABV at varying multiplicities of infection (MOI) for 36 hours (h) and the mRNA level and protein level of TRIM72 were analyzed by qPCR and western blotting. As shown in Fig. 1C-1D, the mRNA and protein levels of TRIM72 were gradually increased along with the RABV infection dose increased. Then mRNA level and protein level of TRIM72 were also analyzed in N2a cells (mouse neuroblastoma cell line) and SK-N-SH cells (human neuroblastoma cell line). As shown in Fig. 1E-1F and Fig. S1A-S1B, the mRNA level and protein level of TRIM72 or human TRIM72 (hTRIM72) gradually increased along with the RABV infection dose increased in N2a cells or SK-N-SH cells. These results indicate that RABV infection induces the up-regulation of TRIM72 in both mouse and neuronal cells.

**Fig. 1.**
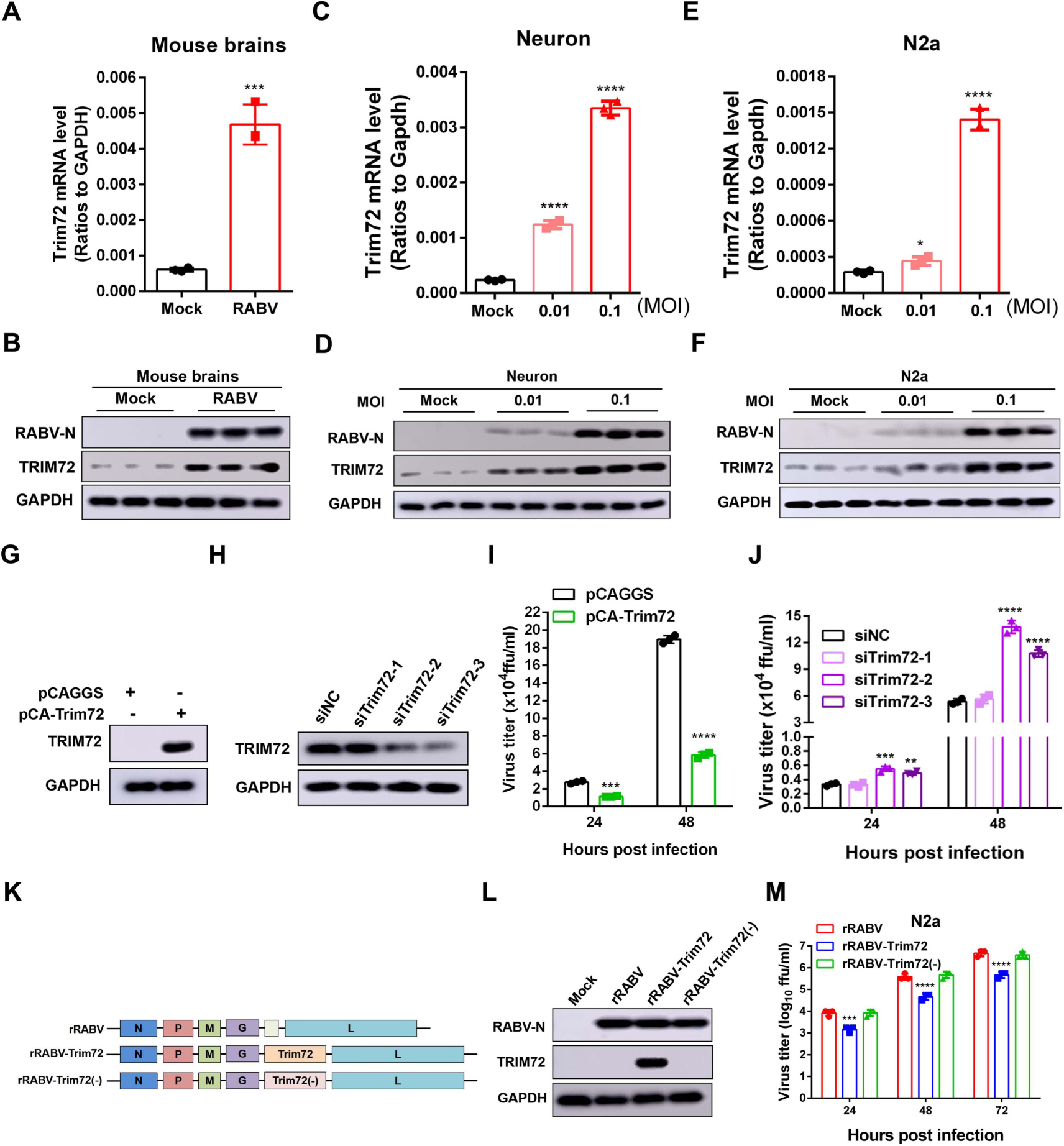
RABV infection induces up-regulation of TRIM72 in mouse brains and neuronal cells, and TRIM72 reduced RABV replication in neuronal cells. (A-B) 6-week-old female C57 BL/6 mice were intracerebrally infected with 200 fluorescence focus units (FFU) RABV (CVS-B2c) or mock-infected. The mouse brains were collected at 6^th^ d post-infection, the mRNA level of TRIM72 was analyzed by qPCR (A), and the protein levels of TRIM72 and RABV-N were analyzed by western blotting (B). (C-D) Primary mouse neuron cells were isolated and cultured, then infected with RABV at different MOI for 36 h. The mRNA level of TRIM72 was analyzed by qPCR (C), and the protein levels of TRIM72 and RABV-N were analyzed by western blotting (D). (E-F) N2a cells were infected with RABV at different MOI for 36 h. The mRNA level of TRIM72 was analyzed by qPCR (E), and the protein levels of TRIM72 and RABV-N were analyzed by western blotting (F). (G) Empty vector (pCAGGS) or TRIM72 over-expression vector (pCA-Trim72) were transfected into N2a cells respectively for 48 h, and then the TRIM72 level was analyzed by western blot. (H) TRIM72-specific siRNAs were transfected into N2a cells respectively for 72 h, then the TRIM72 level was analyzed by western blotting. (I) Empty vector or TRIM72 over-expression vectors were transfected into N2a cells respectively for 12 h, then infected with RABV (MOI=0.01) for the indicated time, and the viral titers in the supernatant were analyzed. (J) TRIM72-specific siRNAs were transfected into N2a cells respectively for 24 h, then infected with RABV (MOI=0.01) for the indicated time, and the viral titers in the supernatant were analyzed. (K) Trim72 and Trim72(−) were inserted into the genome of a recombinant RABV (rRABV), named rRABV-TRIM72 and rRABV-TRIM72(−) respectively (L) N2a cells were infected with different types of rRABVs (MOI=3) for 36 h, then the TRIM72 level was analyzed by western blotting. (M) N2a cells were infected with different types of rRABVs (MOI=0.01) and their growth kinetics were compared. Statistical analysis of grouped comparisons was carried out by student’s t-test (*P < 0.05; **P<0.01; ***P<0.001; ****P<0.0001). The bar graph represents means ± SD, n = 3. Western blot data are representative of at least two independent experiments.

### TRIM72 suppresses RABV replication in neuronal cells

To further investigate the relationship between TRIM72 and RABV, over-expression and silencing systems of TRIM72 were constructed. We then transfected the TRIM72 over-expression vector into N2a cells for 48 h or three pairs of TRIM72 specific small interfering RNA (siTrim72-1, −2, −3) individually into N2a cells for 72 h, and the TRIM72 expression level was analyzed by western blotting. In comparison to the empty vector group, the TRIM72 level was dramatically increased in the TRIM72 over-expression group (Fig. 1G). Conversely, the TRIM72 level was dramatically following transfection with siTRIM72-2 and siTRIM72-3 in N2a cells (Fig. 1H). Next, an empty vector or TRIM72 over-expression vector was transfected into N2a cells for 12 h and then infected the cells with RABV (MOI=0.01). The supernatants of RABV-infected cells were collected at 24 h and 48 h post-infection, then viral titers were analyzed. From the results we can see RABV titer was dramatically decreased post TRIM72 over-expression no matter at 24 h or 48 h post-infection (Fig. 1I). For analysis of the effect of Trim72 silencing on RABV replication, two pairs of siTrim72s (siTRIM72-2 and siTRIM72-3) were transfected into N2a cells individually for 24 h and then the cells were infected with RABV (MOI=0.01), the cell supernatants were collected at 24 h and 48 h post-infection and viral titers were analyzed. From the results, we can see that RABV titer dramatically increased post TRIM72 silencing both at 24 h and 48 h post-infection (Fig. 1J). Moreover, we found that human TRIM72 (hTRIM72) over-expression dramatically reduced the RABV titer in SK-N-SH cells (Fig. S1C-S1D). To further analyze the function of TRIM72 in restricting RABV, the recombinant RABVs (rRABVs) were constructed. The recombinant viruses were derived from the CVS-B2c strain, including unaltered rRABV, rRABV harboring TRIM72 coding sequence (rRABV-Trim72) and rRABV harboring the TRIM72 sequence which ATG mutation to CTG (rRABV-Trim72(−)) (Fig. 1K). N2a cells were infected with those rRABVs (MOI=3) for 36 h, then the TRIM72 expression level was analyzed by western blotting. As expected, the TRIM72 protein level was extremely higher in rRABV-TRIM72 infected N2a cells than in rRABV or rRABV-TRIM72(−) infected cells (Fig. 1L). Subsequently, virus growth kinetics experiments were performed in N2a cells, and N2a cells were infected with the rRABVs (MOI=0.01) separately, the supernatants were collected at the indicated time points post rRABVs infection for analysis of virus titers. The virus titer was significantly lower in the rRABV-TRIM72 infected cells than in rRABV- and rRABV-TRIM72(−)-infected cells (Fig. 1M). Above all, over-expression of TRIM72 reduced the viral titer of RABV in neuronal cells.

### TRIM72 reduces RABV pathogenicity *in vivo*

To investigate the role of TRIM72 in RABV infection in vivo, the pathogenicity of rRABV, rRABV-Trim72 and rRABV-Trim72(−) in the C57 BL/6 mouse model was compared. Mice were infected intranasally with rRABV, rRABV-Trim72, or rRABV-Trim72(−) at a dose of 100 FFU/mouse. Then the body weight, clinical score and survival were monitored. The body weight of the mice infected with rRABV or rRABV-Trim72(−) decreased dramatically from day 7 to day 13 post-infection (p.i.), whereas the body weight of those mice infected with rRABV-Trim72 showed a slight decrease from day 13 to day 17 p.i. (Fig. 2A). The rabies clinical symptoms of rRABV- or rRABV-Trim72(−) -infected mice were appeared from day 6 to day 13, whereas the mice infected with rRABV-TRIM72 display mild symptoms from day 11 to day 17 (Fig. 2B). Finally, 80% of the mice infected with rRABV-Trim72 survived, and only 10% and 20% survival ratio for rRABV- and rRABV-Trim72(−) -infected mice, respectively (Fig. 2C).

**Fig. 2.**
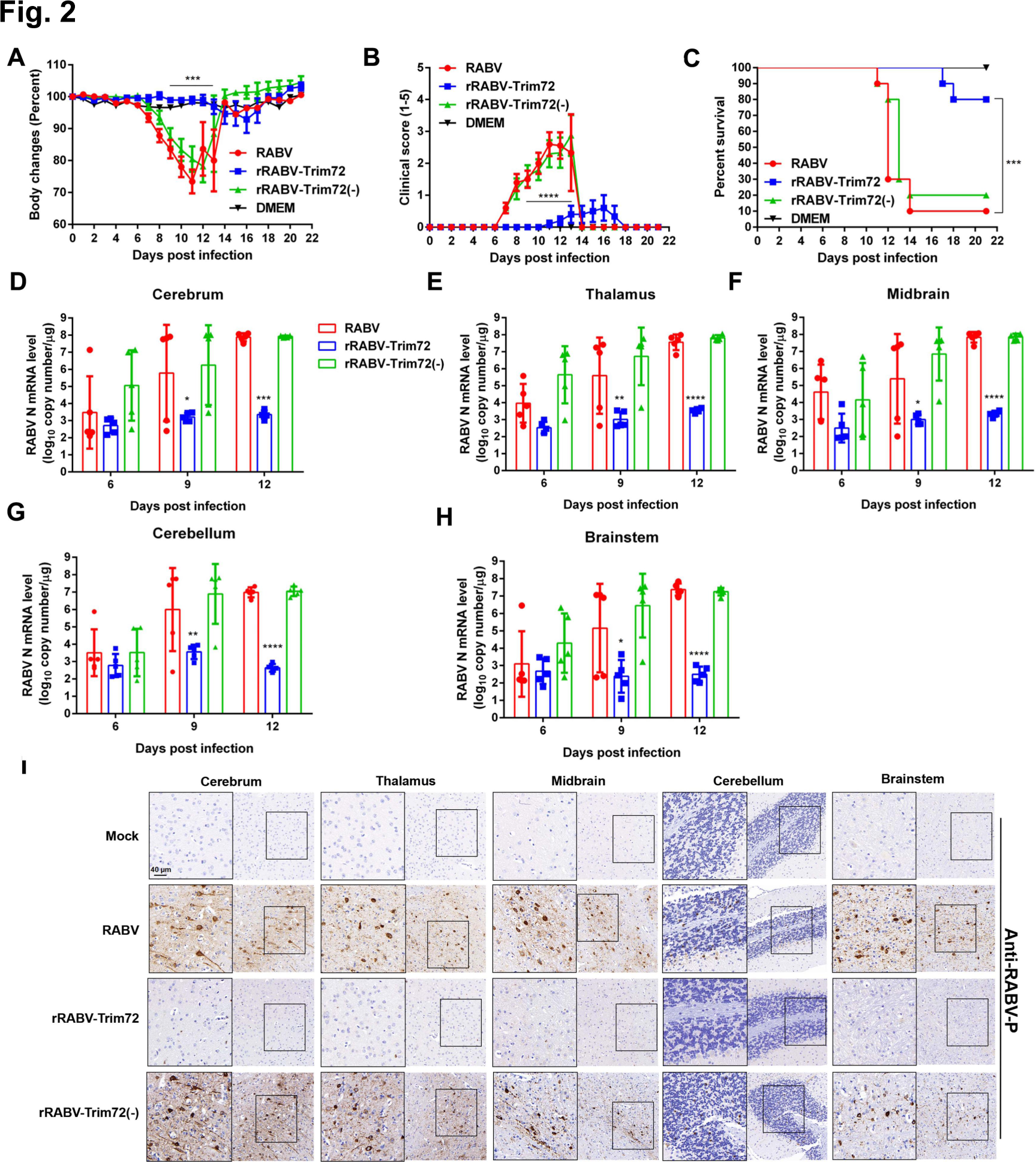
TRIM72 attenuates RABV pathogenicity in vivo. (A-C) Female C57 BL/6 mice (8-week-old, n=10) were intranasally infected with 100 FFU rRABV, rRABV-Trim72, rRABV-Trim72(−), or were mock-infected. Body weight change (A), clinical score (B), and survival ratio (C) were monitored daily for continuous 3 weeks. (means ± SEM; ***P<0.001; ***P<0.0001; body weight change and the clinical score were analyzed by Two-way ANOVA test; survival ratio was analyzed by log-rank test). (D-H) At indicated time points, different brain parts of the infected mice were collected to analyze the levels of RABV N mRNA by qPCR. (n=5; means ± SEM; *P < 0.05; **P<0.01; ***P<0.001; ***P<0.0001 by student’s Two-way ANOVA test). (I) At the 12th day post-infection, mouse brains were collected, sectioned, and analyzed by immunohistochemistry using antibodies against RABV P. Scale bar, 40 μm.

To analyze the viral loads in the brains of mice infected with rRABV, rRABV-Trim72 and rRABV-Trim72(−), the RABV viral N mRNA levels were analyzed in different brain parts post that virus infection via intranasally (100 FFU/mouse). On day 6, day 9 and day12 p.i., the brains infected with different viruses were collected and divided into 5 parts (Cerebrum, Thalamus, Midbrain, Cerebellum and Brainstem), then the N mRNA levels in different brain parts were analyzed by qPCR. Dramatically reduced RABV N mRNA levels in rRABV-Trim72-infected different mice brain parts were presented compared with rRABV- or rRABV-Trim72(−)-infected mouse brains at day 9 and day 12 p.i. (Fig. 2D-2H). Then the immunohistochemistry assay was performed by using an anti-RABV-P antibody to analyze the RABV protein level in different brain parts post virus infection for 12 days. Almost no RABV-P antigen was observed in rRABV-Trim72 infected brain parts, whereas there displays comparable level of RABV-P antigen in rRABV- or rRABV-Trim72(−)-infected mice brain parts (Fig. 2I). Above all, over-expression of TRIM72 dramatically reduced the pathogenicity of RABV in mice.

### TRIM72 restricts RABV release

Our results demonstrate that TRIM72 inhibits RABV in vitro and in vivo, while the mechanism remains unclear. Therefore, the effects of TRIM72 over-expression on the different stages of the RABV life cycle were analyzed. First, the TRIM72 over-expression vector or an empty vector was individually transfected into N2a cells for 36 h and then the cells were infected with RABV (MOI=0.1) for 1 h at 4℃, the cells were collected and the viral genomic RNA (vRNA) level was analyzed by qPCR to evaluate the effect of TRIM72 on RABV attachment. As shown in Fig. 3A, there is no obvious difference in viral genomic RNA levels between the control group and the TRIM72 over-expression group, indicating that TRIM72 did not affect RABV attachment. Next, the effect of TRIM72 over-expression on RABV entry was analyzed, TRIM72 over-expression vector or empty vector was individually transfected into N2a cells for 36 h and then infected with RABV (MOI=0.1) for 2 h at 37℃, the cells were collected and the vRNA levels were analyzed by qPCR. There is no obvious difference in vRNA levels between the control group and the TRIM72 over-expression group (Fig. 3B), indicating that TRIM72 did not affect RABV entry. Then TRIM72 over-expression vector or empty vector was individually transfected into N2a cells for 12 h and then RABV was infected (MOI=0.01), the cells were collected at 24 h and 48 h post-infection, and the RABV N mRNA level and vRNA level were analyzed by qPCR. As shown in Fig. 3C-3D, there is almost no difference in RABV N mRNA level and vRNA level between the TRIM72 over-expression group and the empty vector group. These results indicate that TRIM72 over-expression does not affect RABV transcription and replication. Then virus-spreading assay was performed to analyze the impact of TRIM72 on RABV spreading between cells. N2a cells were infected with rRABV, rRABV-Trim72, or rRABV-Trim72(−) individually (MOI=0.001), then the cell culture medium containing 1% agar was added to cover the N2a cells for 48 h. Fluorescence in isothiocyanate (FITC)-conjugated anti-RABV-P protein antibody (FITC-P) was stained post removed from the cell culture medium and the fluorescence plaque was analyzed. As shown in Fig. 3E, there is almost no difference in the size of fluorescence plaque and the number of infected cells in these N2a cells (Fig. 3E-3F), indicating that TRIM72 does not affect RABV spreading between cells.

**Fig. 3.**
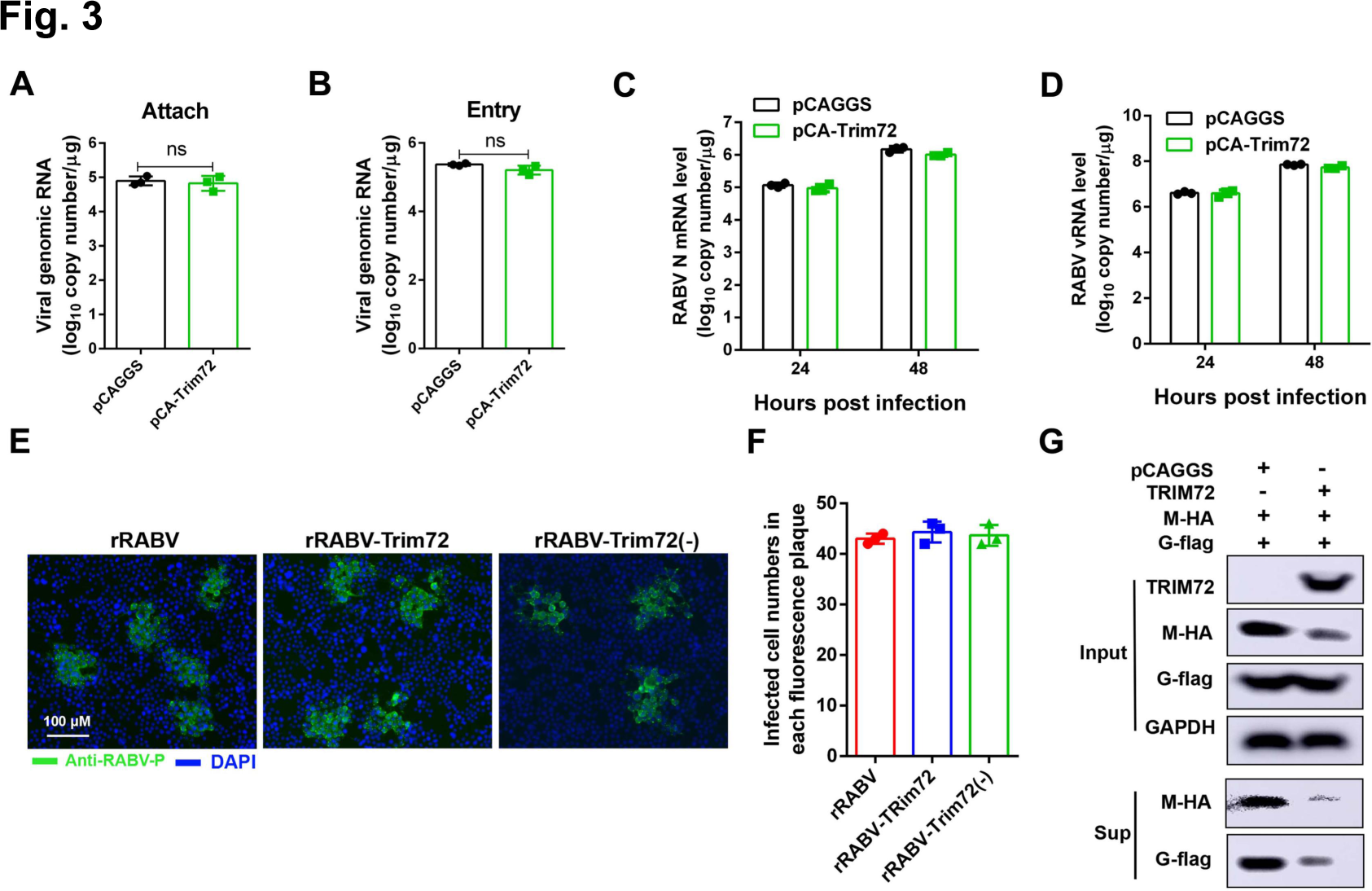
TRIM72 restricts RABV infection at the viral release stage. (A) TRIM72 over-expression vector or empty vector were transfected into N2a cells respectively for 36 h, then infected with RABV (MOI=0.1) for 1 h at 4℃, the cells were collected and the viral genomic RNA (vRNA) levels were analyzed by qPCR. (B) TRIM72 over-expression vector or empty vector was transfected into N2a cells individually for 36 h then infected with RABV (MOI=0.1) for 2 h at 37℃, the cells were collected and the vRNA levels were analyzed by qPCR. (C-D) TRIM72 over-expression vector or empty vector were transfected into N2a cells individually for 12 h and then infected with RABV (MOI=0.01) for the indicated time points, the cells were collected and the levels of RABV-N-mRNA (D) and vRNA (E) were analyzed by qPCR. (E-F) N2a cells were infected with rRABV, rRABV-Trim72, or rRABV-Trim72(−) at MOI 0.005. Then the cells were covered with culture medium containing 1% agar and incubated for 48 h. The viral spread was compared and the infected cell numbers were calculated within the fluorescence focus. Scale bar, 100 μm. (G) pCAGGS or pCA-Trim72 together with pCA-M and pCA-G were co-transfected into HEK-293T cells respectively for 48 h, and the protein levels of M and G in the supernatants and cells were analyzed by western blotting. Western blot data are representative of at least two independent experiments.

Our results clearly show that TRIM72 over-expression dramatically reduced the RABV titer in N2a cells (Fig. 1I and Fig. S1D). However, TRIM72 over-expression almost does not affect RABV attachment, entry, transcription, replication and spreading in N2a cells. Therefore, we hypothesized that TRIM72 might affect RABV release. To investigate the hypothesis, a virus-like particle (VLP) system was constructed as previously reported to analyze the effect of TRIM72 over-expression on RABV release [28, 29]. TRIM72 over-expression vector or empty vector together with RABV-M and RABV glycoprotein (G) were co-transfected into 293T cells for 48 h and the VLP production in the supernatants was analyzed using western blotting. As shown in Fig. 3G, the protein levels of M and G in the supernatants were dramatically decreased post TRIM72 over-expression compared to the control group, indicating that TRIM72 restricts VLP production. As our speculation, these results confirm our hypothesis that TRIM72 over-expression restricts RABV release.

### TRIM72 induces proteasomal degradation of RABV-M by promoting its K48-linked ubiquitination

Our results revealed that TRIM72 restricts RABV release. However, an interesting observation was the significant decrease in M protein levels following TRIM72 over-expression (Fig. 3G). Given that TRIM72 is an E3 ubiquitin ligase, we speculated that its inhibitory effect on RABV may be responsible for its E3 ubiquitin ligase function, leading to the degradation of the viral proteins. To explore this hypothesis, over-expression vectors of RABV viral proteins (N, P, M, G), together with TRIM72 expression vector or an empty vector, were individually co-transfected into N2a cells for 48 h, the cells were collected and the viral protein levels were analyzed by western blotting. As shown in Fig. 4A, the protein levels of RABV-N, -P, or -G almost have no changes following TRIM72 over-expression. In contrast, the protein level of RABV-M was dramatically decreased in the TRIM72 over-expression group compared to the empty vector group, indicating that TRIM72 specifically induces the degradation of RABV-M. We also found that hTRIM72 can induce the degradation of RABV-M in SK-N-SH cells (Fig. S2), indicating the conserved function of TRIM72 in degrading RABV-M. Furthermore, proteasome inhibitor Mg132 and lysosome inhibitor NH_4_Cl were used individually in N2a cells to elucidate the degradation pathway of M induced by TRIM72. As shown in Fig. 4B, TRIM72 over-expression remains induces the degradation of M even after NH_4_Cl treatment, whereas the degradation was interrupted following Mg132 treatment. Then a dose-dependent assay was performed, and the M protein level was gradually decreased along with increasing TRIM72 level (Fig. 4C). However, the degradation was impeded after Mg132 treatment (Fig. 4D). These findings indicate that TRIM72 induces RABV-M degradation through the proteasome pathway.

**Fig. 4.**
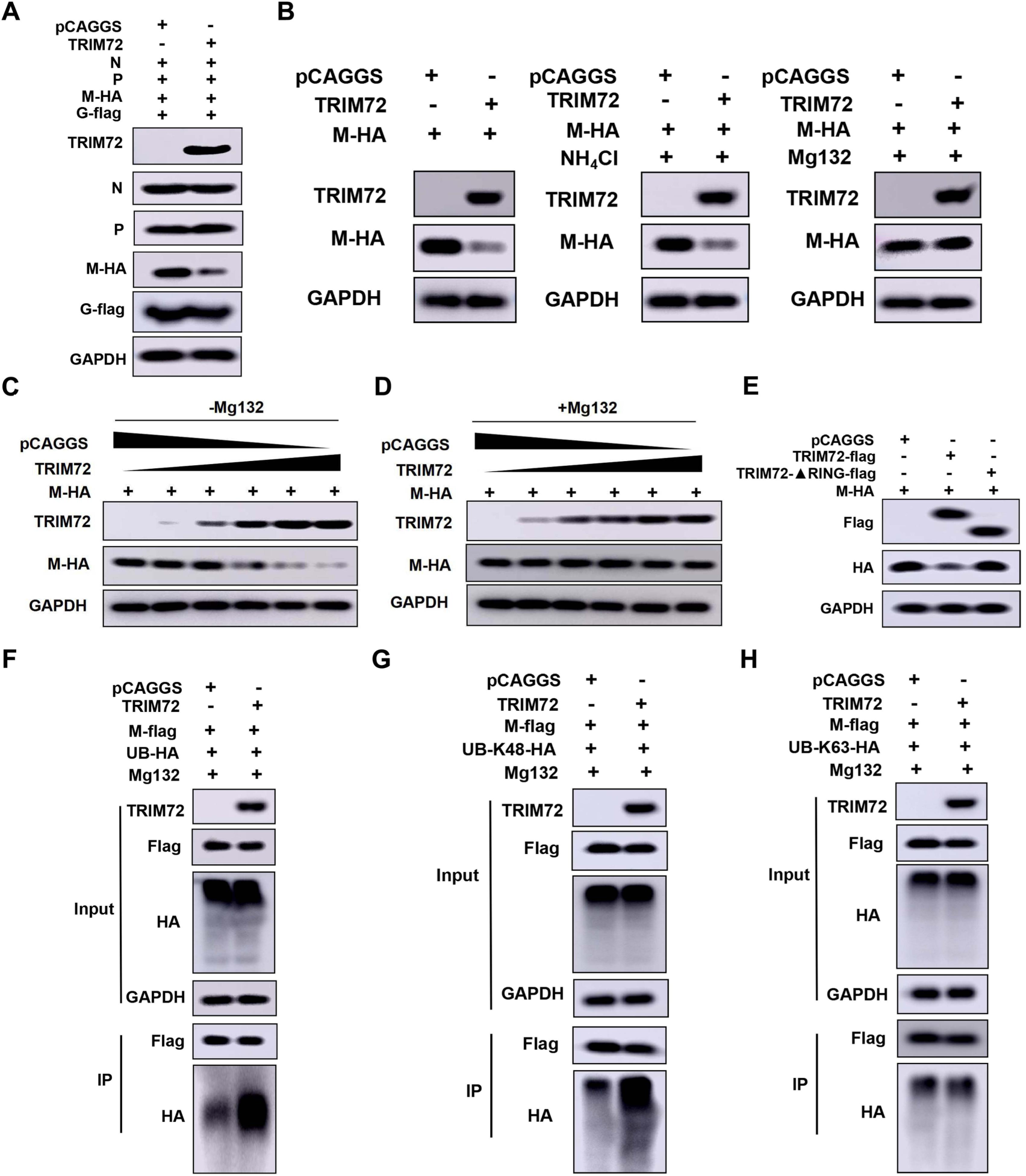
TRIM72 induces proteasomal degradation of RABV-M by promoting K48-linked ubiquitination. (A) pCAGGS or pCA-Trim72 together with pCA-N, pCA-P, pCA-M, and pCA-G were co-transfected into N2a cells respectively for 48 h, and the protein levels of N, P, M and G were analyzed by western blotting. (B) pCAGGS or pCA-Trim72 together with pCA-M-HA were co-transfected into N2a cells respectively. The specific inhibitors for proteasome and lysosome, Mg132 (10 µM) and NH_4_Cl (5 mM) were applied. Then the protein levels of TRIM72 and M were analyzed by western blotting. (C) pCA-M-HA together with pCAGGS or different concentrations of pCA-Trim72 were co-transfected into N2a cells for 48 h. Then the protein levels of TRIM72 and M were analyzed by western blotting. (D) pCA-M-HA together with pCAGGS or different concentrations of pCA-Trim72 were co-transfected into N2a cells. Then Mg132 was applied and the protein levels of TRIM72 and M were analyzed by western blotting. (E) pCA-M-HA together with pCAGGS, pCA-Trim72-flag, or Trim72-ΔRING-flag were co-transfected into N2a cells respectively for 48 h, then M-HA level was analyzed by western blotting. (F) pCAGGS or pCA-Trim72 together with pCA-M-flag and UB-HA over-expression vectors were co-transfected into N2a cells respectively. Then treated with Mg132 and Co-IP assays were performed with anti-flag antibody. The protein levels of TRIM72, M-flag, and UB-HA were analyzed by western blotting. (G) pCAGGS or pCA-Trim72 together with pCA-M-flag and UB-K48-HA over-expression vectors were co-transfected into N2a cells respectively. Then treated with Mg132 and Co-IP assays were performed with anti-flag antibody post transfection for 48 h. Then the protein levels of TRIM72, M-flag, and UB-K48-HA were analyzed by western blotting. (H) pCAGGS or pCA-Trim72 together with pCA-M-flag and UB-K63-HA over-expression vector were co-transfected into N2a cells respectively. Then treated with Mg132 and Co-IP assays were performed with anti-flag antibody. The protein levels of TRIM72, M-flag, and UB-K63-HA were analyzed by western blotting. Western blot data are representative of at least two independent experiments.

We then speculated that TRIM72-induced degradation of M via the proteasome pathway was due to its E3 ubiquitin ligase function which promotes M ubiquitination. The RING domain is essential for TRIM72 to promote the ubiquitination of the target proteins [25]. Thus, TRIM72, a RING domain deficient TRIM72 (TRIM72-ΔRING) together with M were co-overexpressed in N2a cells for 48 h and the M level was analyzed by western blotting. As shown in Fig. 4E, WT TRIM72 over-expression induces M degradation, whereas TRIM72ΔRING over-expression failed to induce M degradation. These results indicate that TRIM72 induces M degradation depending on its RING domain. We then examined the effect of TRIM72 on the ubiquitination of M. The ubiquitination level of M was dramatically increased following TRIM72 over-expression in the presence of Mg132 (Fig. 4F), indicating that TRIM72 promotes the ubiquitination of M. K48- and K63-linked polyubiquitin chains are the two major types of ubiquitin linkages [30], then we further characterized which type of ubiquitination of M was modified by TRIM72. TRIM72 over-expression plasmids and M over-expression plasmids together with either K48- or K63-ubiquitin over-expression plasmids were co-transfected into N2a cells for 48 h respectively, then ubiquitination assays were performed. As shown in Fig. 4G-4H, TRIM72 specifically promotes the K48-linked ubiquitination of M, while almost does not affect the K63-linked ubiquitination of M. Overall, TRIM72 promotes K48-linked ubiquitination of M, consequently facilitating M degradation via the proteasome pathway.

### The SPRY domain of TRIM72 directly interacts with RABV-M

Our above results demonstrate that TRIM72 promotes the K48-linked ubiquitination of RABV-M, thus we speculated that TRIM72 might interact with RABV-M directly. Consequently, Co-Immunoprecipitation (Co-IP) assays were performed to investigate this interaction, as shown in Fig. 5A-5B and Fig. S3, both murine TRIM72 and hTRIM72 were found to interact directly with M. For further analysis of the interaction between TRIM72 and RABV-M, different TRIM72 truncation over-expression vectors were designed and constructed based on its functional domain (Fig. 5C). These vectors together with M over-expression vectors were co-overexpressed in N2a cells for 48 h and Co-IP assays were performed. As shown in Fig. 5D, the TRIM72 truncations which contain the SPRY domain could interact with M, indicating that the TRIM72-SPRY domain directly interacts with M. Additionally, different M truncation over-expression vectors were also constructed based on its secondary structure (Fig. 5E). Co-IP assays were performed between M truncations and TRIM72-SPRY domain. As shown in Fig. 5F, Both the M truncations were capable of interacting with the TRIM72-SPRY domain, indicating that M truncation (49-160 aa) was the key interaction domain that interacts with the TRIM72-SPRY domain.

**Fig. 5.**
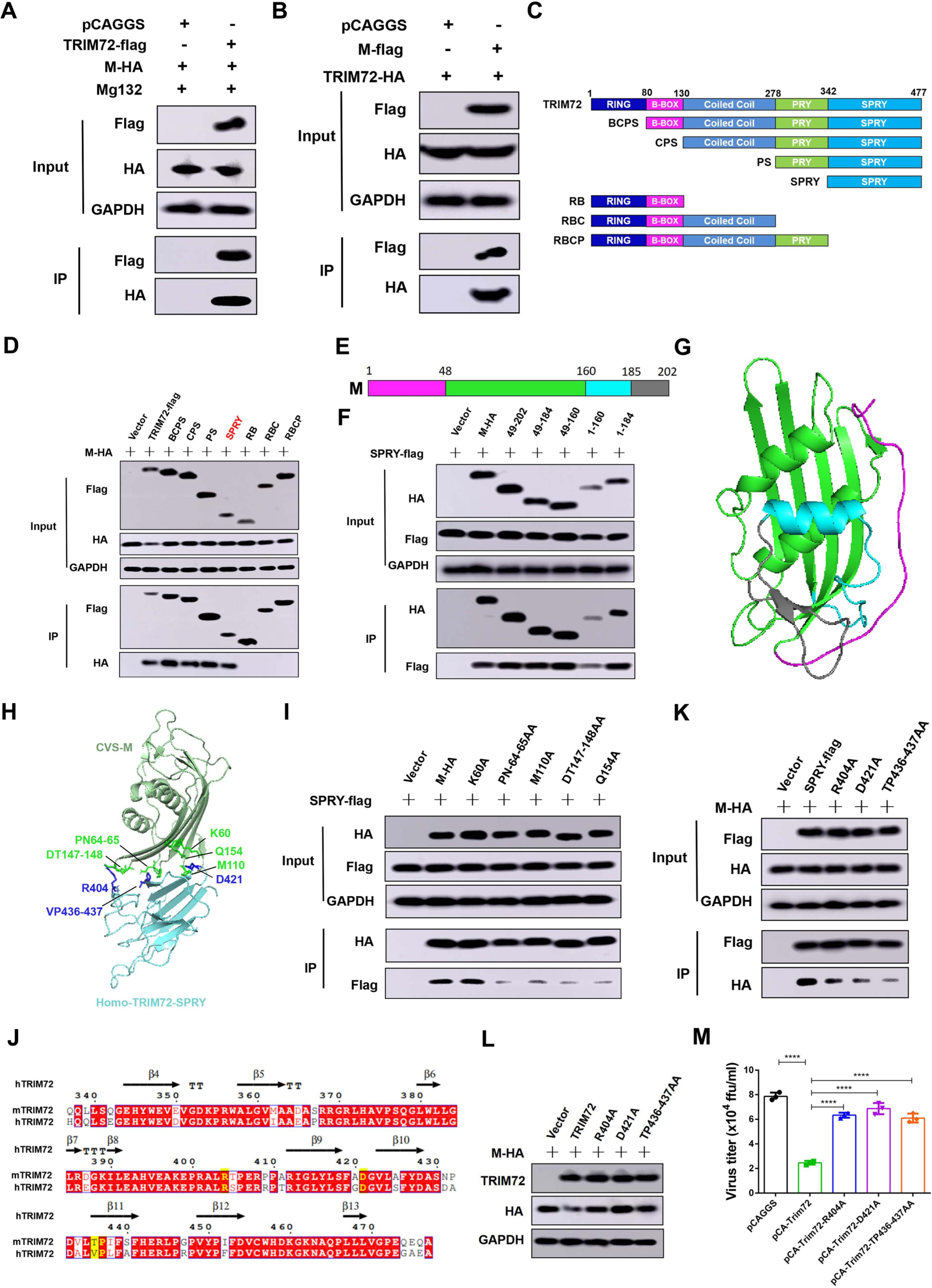
TRIM72 directly interacts with RABV-M via the SPRY domain. (A) pCAGGS or pCA-Trim72-flag together with pCA-M-HA were co-transfected into N2a cells respectively. Then Mg132 (10 µM) was treated, and Co-IP assays were performed with anti-flag antibody post-transfection for 48 h. The protein levels of TRIM72-flag and M-HA were analyzed by western blotting. (B) pCAGGS or pCA-M-flag together with pCA-Trim72-HA were co-transfected into N2a cells respectively for 48 h. Then Co-IP assays were performed with anti-flag antibody. The protein levels of TRIM72-flag and M-HA were analyzed by western blotting. (C) TRIM72 truncations were designed and constructed based on its functional domain. (D) The over-expression vectors of TRIM72 truncations together with pCA-M-HA were co-transfected into N2a cells respectively for 48 h. Then Co-IP assays were performed with anti-flag antibody. The protein levels of TRIM72 truncations and M-HA were analyzed by western blotting. (E) RABV-M truncations were designed and constructed based on its secondary structure. (F) The over-expression vectors of M truncations together with pCA-Trim72-SPRY-flag were co-transfected into N2a cells for 48 h. Then Co-IP assays were performed with anti-HA antibody. The protein levels of M truncations and SPRY-flag were analyzed by western blotting. (G) A structural model of RABV-M was built using SWISS-MODEL online software (https://swissmodel.expasy.org/interactive) based on the crystal structure of Lagos bat virus M protein (PDB code: 2W2S). (H) An interaction model of the human TRIM72-SPRY domain and RABV-M was built with GalaxyWEB online software (https://galaxy.seoklab.org/) based on hTRIM72 structure (PDB code: 6NPY), the potential interaction sites were labeled. (I) Over-expression vectors of RABV-M truncations together with TRIM72-SPRY over-expression vector were co-transfected into N2a cells respectively for 48 h. Then Co-IP assays were performed with anti-HA antibody. The protein levels were analyzed by western blotting. (J) The protein sequences of hTRIM72 and mouse TRIM72 were compared and analyzed with ESPript 3.0 online software (https://espript.ibcp.fr/ESPript/cgi-bin/ESPript.cgi). (K) Over-expression vectors of TRIM72-SPRY mutations together with M-HA over-expression vectors were co-transfected into N2a cells respectively for 48 h. Then Co-IP assays were performed with anti-flag antibody and protein levels were analyzed by western blotting. (L) Over-expression vectors of TRIM72 mutations together with M-HA over-expression vector were co-transfected into N2a cells respectively for 48 h. The protein levels were analyzed by western blotting. (M) Over-expression vectors of TRIM72 mutations or empty vectors were transfected into N2a cells respectively for 12 h, then infected with rRABV (MOI=0.01) for 48 h and the viral titers in the supernatants were analyzed. Statistical analysis of grouped comparisons was carried out by student’s t-test (*P < 0.05; **P<0.01; ***P<0.001; ****P<0.0001). The bar graph represents means ± SD, n = 3. Western blot data are representative of at least two independent experiments.

We then attempted to construct an M-TRIM72-SPRY interaction model to analyze the specific interaction sites. Science there was no available structure model for RABV-M, thus we built a RABV-M model based on the Lagos bat virus M protein structure with SWISS-MODEL online software. The RABV-M model is shown in Fig. 5G. A previous study has reported the hTRIM72 crystal structure, allowing us to employ it as the model for building the M-TRIM72-SPRY interaction model using GalaxyWEB online software. The M-TRIM72-SPRY interaction model was shown in Fig. 5H, and the potential interaction sites between M and TRIM72-SPRY were analyzed and presented. We found that the residues R404, D421 and VP436-437 of hTRIM72-SPRY could interact with the residues K60, PN64-65, M110, DT147-148 and Q154 of M respectively. The potential interaction residues were then validated by mutation and Co-IP assays. As shown in Fig. 5I, the protein level of SPRY that was pulled down by mutated M (PN64-65AA, M110A, DT147-148AA and Q154A) was dramatically decreased compared to that of WT M. Since the amino acid sequence homology between hTRIM72-SPRY and murine TRIM72-SPRY reached 84.3% (Fig. 5J). Therefore, the corresponding interaction residues of the murine TRIM72-SPRY domain were mutated and Co-IP experiments were performed to determine the changes in binding between the mutated murine TRIM72-SPRY and M. As shown in Fig. 5K, the M protein level pulled down by mutated TRIM72-SPRY (R404A, D421A and VP436-437AA) dramatically reduced compared to that of WT TRIM72-SPRY. In conclusion, these results strongly support the reliability of the key interaction sites between M and TRIM72-SPRY as analyzed by the predicted M-TRIM72-SPRY interaction model. Specifically, residues PN64-65, M110, DT147-148 and Q154 on M, as well as R404, D421, and VP436-437 on TRIM72-SPRY, have been identified as critical for the interaction between TRIM72-SPRY and M.

Furthermore, we sought to analyze whether these interaction site mutations in TRIM72-SPRY disrupted the function of TRIM72 in M degradation, then the over-expression vectors of full-length TRIM72 mutations (R404A, D421A and VP436-437AA) were constructed and together with M over-expression vector were co-transfected into N2a cells for 48 h. Then the M levels were analyzed by western blotting. Interestingly, the protein level of M was comparable in the TRIM72 mutated group compared to the empty vector group, while the M level decreased significantly following WT TRIM72 over-expression (Fig. 5L). Moreover, the antiviral function of TRIM72 mutations was also analyzed. The over-expression vectors of WT full-length TRIM72 or mutations were transfected into N2a cells for 12 h and then infected with RABV (MOI=0.01) for 48h, the virus titers in the cell culture supernatant were measured. As shown in Fig. 5M, the antiviral function of TRIM72 was impaired following R404, D421 and VP436-437 mutation in TRIM72 compared to the WT TRIM72 group.

### TRIM72 promotes the ubiquitination of RABV-M at the K195 site

To identify the specific lysine site in M that is ubiquitinated by TRIM72, the lysine in M was mutated respectively (Fig. 6A). The over-expression vectors of M which lysine (K) mutated to alanine (A) were constructed and together with TRIM72 were co-overexpressed in N2a cells for 48 h respectively, then western blots were performed and the protein level of mutated M was analyzed. As shown in Fig. 6B, only the M-K195A mutant abolished the degradation of M which was induced by TRIM72, indicating that the K195 is a potential lysine residue targeted by TRIM72 for M ubiquitination. Subsequently, the stability of WT M and M-K195A were compared when individually over-expressed in N2a cells for 48 h, the M level was analyzed by western blot. As shown in Fig. 6C, The M-K195A protein level was higher than that of WT M, indicating that M-K195A was much more stable than WT M. Then we examined the ubiquitination level of M-K195A which was induced by TRIM72. Empty vector or TRIM72 over-expression vector together with ubiquitin and M-K195A were co-overexpressed in N2a cells for 48 h and then the ubiquitination level of M-K195A was analyzed. As shown in Fig. 6D, M-K195A presents a comparable ubiquitination level between the vector group and TRIM72 over-expression group, demonstrating that the K195A mutation abolished the enhanced ubiquitination level induced by TRIM72. These results strongly indicate that the K195 site is a critical ubiquitination target for M, mediated by TRIM72.

**Fig. 6.**
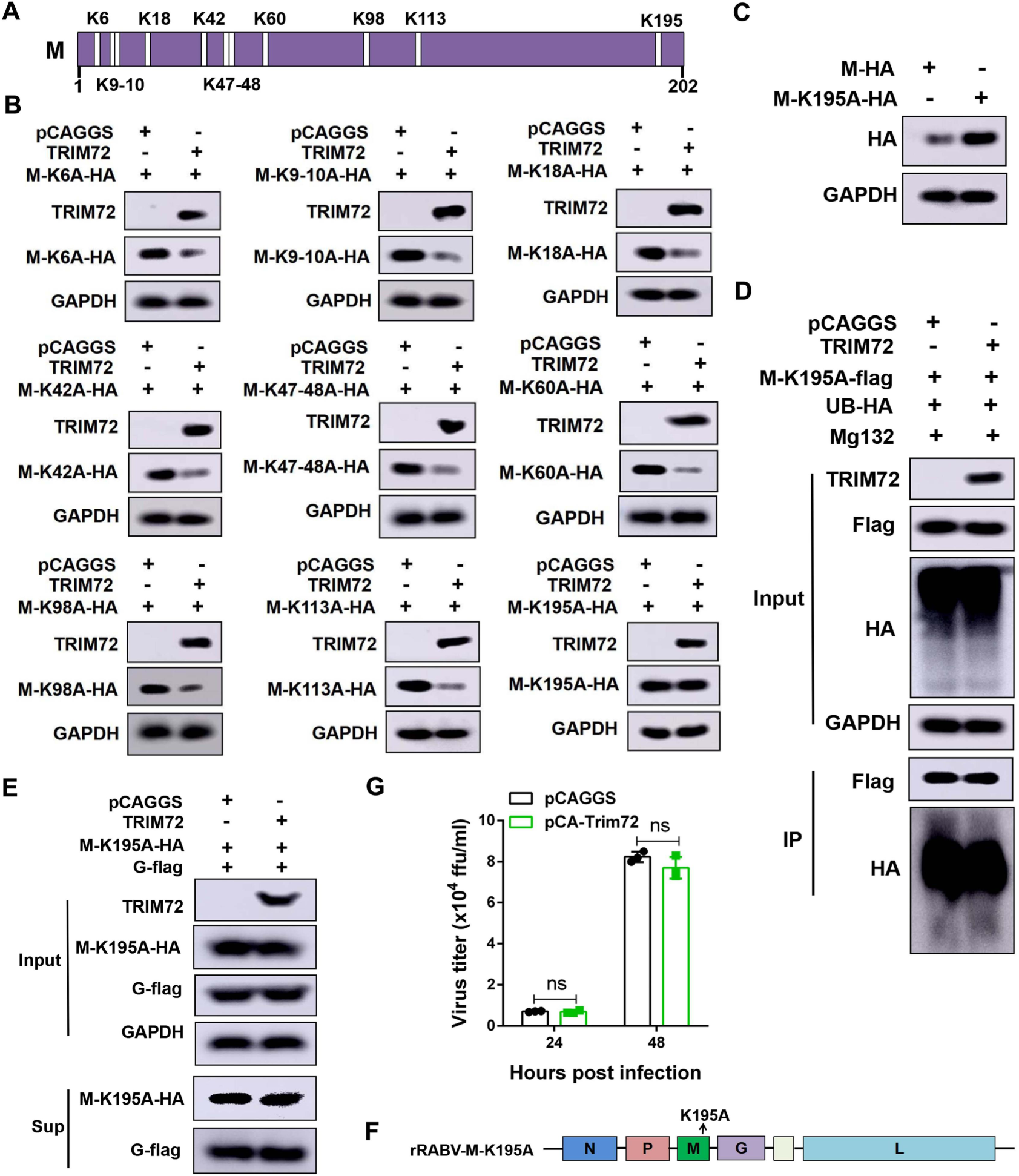
TRIM72 induces ubiquitination of RABV-M at K195. (A) Illustration of the lysine (K) sites in RABV-M. (B) The lysine residues in RABV-M were mutated to alanine and over-expression vectors were constructed and together with pCAGGS or pCA-Trim72 were co-transfected into N2a cells respectively for 48 h. The protein levels of M mutations and TRIM72 were analyzed by western blotting. (C) WT M-HA or M-K195A was over-expressed in N2a cells respectively for 48 h, and the protein level was analyzed by western blotting. (D) pCA-M-K195A-flag together with pCAGGS or pCA-Trim72 were co-transfected into N2a cells and treated with Mg132 (10 µM), then Co-IP assays were performed with anti-flag antibody. The protein levels were analyzed by western blotting. (E) pCAGGS or pCA-Trim72 together with pCA-M-K195A and pCA-G were co-transfected into HEK-293T cells respectively for 48 h, the protein levels of M-K195A and G in the supernatants and cells were analyzed by western blotting. (F) Schematic diagram of the recombinant rRABV-M-K195A. (G) pCAGGS or pCA-Trim72 was transfected into N2a cells respectively for 12 h, then infected with rRABV-M-K195A (MOI=0.01) for 48 h. The supernatants were collected and viral titers were analyzed. Western blot data are representative of at least two independent experiments.

To further investigate the impact of the M-K195A mutation on the effect of TRIM72 on RABV release, the release assay was performed. TRIM72 or empty vector together with M-K195A and RABV G were co-overexpressed in 293T cells for 48 h and the VLP production in the supernatants was analyzed by western blotting. As shown in Fig. 6E, the protein levels of M-K195A and G in the supernatants were comparable between the TRIM72 over-expression group and the control group, indicating that TRIM72 lost its ability to restrict the VLP production following M-K195 site mutation. Moreover, A M-K195A mutant RABV (rRABV-M-K195A) was constructed (Fig. 6F), and the restriction effect of TRIM72 on rRABV-M-K195 was analyzed. N2a cells were transfected with either a vector or TRIM72 over-expression vector for 12 h and then infected with rRABV-M-K195A (MOI=0.01). The supernatants were collected and viral titers were analyzed at 48 h post-infection. As shown in Fig. 6G, there are almost no obvious changes in viral titers between the vector and the TRIM72 over-expression group. These results indicate that TRIM72 restricts RABV release by promoting the ubiquitination of M at the 195 site, thereby promoting the degradation of the M protein.

### TRIM72 targeting the K195 site is partially conserved among lyssavirus M

RABV is a member of lyssaviruses, the M proteins display a high homology among lyssaviruses (Fig. S4). Therefore, we speculated that TRIM72 might also induce the degradation of other lyssavirus M proteins. Our results demonstrate that TRIM72 promotes CVS-M-K195 ubiquitination thereby promoting the degradation of CVS-M protein. Then the K195 site was analyzed among the M proteins of those lyssaviruses. As shown in Fig. S4, the K195 site was partially conserved among those lyssavirus M proteins, such as dog-derived RABV M (named DRV-M), ABLV-M (At the 205 site in ABLV-M), DUVV-M and EBLV-M, while the 195 site was the E instead of K in LBV-M or MOKV-M. Firstly, we tested the effect of TRIM72 on DRV-M, the DRV-M together with TRIM72 or empty vector were co-overexpressed in N2a cells for 48 h, and then western-blot was performed to analyze the DRV-M protein level. As shown in Fig. 7A, the DRV-M protein level decreased significantly following TRIM72 over-expression, indicating that TRIM72 induce the degradation of DRV-M. The other M proteins containing the K195 site of these lyssaviruses were also tested including ABLV-M, DUVV-M and EBLV-M. As shown in Fig. 7B-7D, similar results were observed, where the protein levels of ABLV-M, DUVV-M and EBLV-M were dramatically reduced following TRIM72 overexpression, indicating that TRIM72 induced the degradation of those lyssavirus M proteins. Importantly, treatment with Mg132, a proteasome inhibitor, could reverse the degradation of lyssavirus M proteins induced by TRIM72 over-expression (Fig. 7E-7H). Moreover, the results of Co-IP assays demonstrated that DRV-M, ABLV-M, DUVV-M and EBLV-M could also interact with TRIM72 (Fig. 7E-7H). However, when we tested the effect of TRIM72 on LBV-M and MOKV-M, we found that although TRIM72 could interact with LBV-M or MOKV-M, TRIM72 did not induce the degradation of LBV-M or MOKV-M (Fig. S5).

**Fig. 7.**
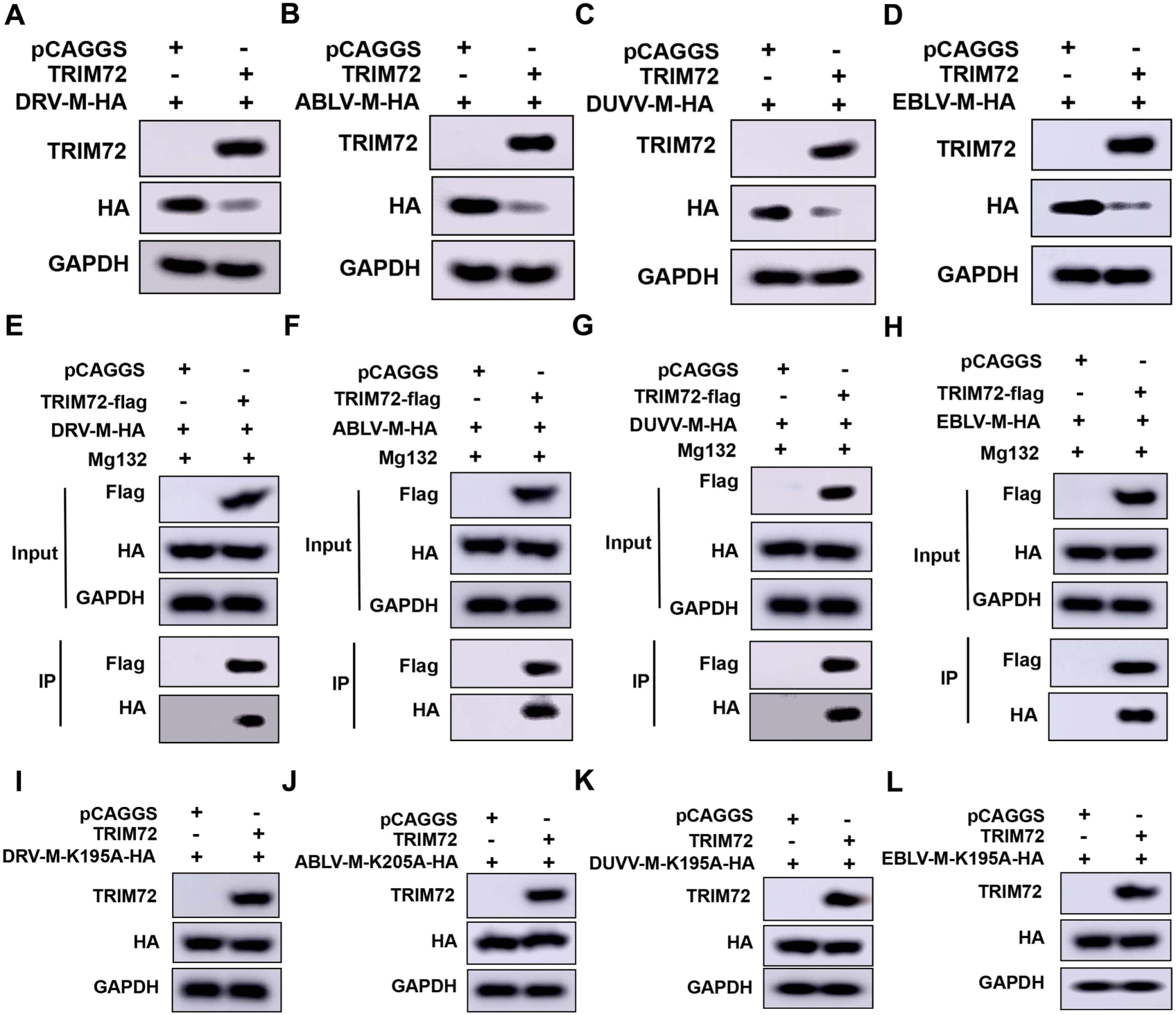
K195 is the conserved ubiquitination site among lyssavirus targeted by TRIM72. (A-D) HA-tagged lyssavirus M proteins (DRV-M-HA, ABLV-M-HA, DUVV-M-HA and EBLV-M-HA) together with empty vector or TRIM72 were co-overexpressed in N2a cells for 48 h, then protein levels of DRV-M-HA (A), ABLV-M-HA (B), DUVV-M-HA (C) and EBLV-M-HA (D) were analyzed by western blotting. (E-H) HA-tagged lyssavirus M proteins (DRV-M-HA, ABLV-M-HA, DUVV-M-HA, and EBLV-M-HA) together with empty vector or TRIM72-flag were co-overexpressed in N2a cells respectively, followed by treatment with Mg132 (10 µM). Co-IP assays were performed with anti-flag antibody post-transfection for 48 h, and protein levels of DRV-M-HA (E), ABLV-M-HA (F), DUVV-M-HA (G), and EBLV-M-HA (H) were analyzed by western blotting. (I-L) The K195 sites (K205 site in ABLV-M) in lyssavirus M proteins were mutated to Alanine, then together with empty vector or TRIM72 were co-overexpressed in N2a cells for 48 h. The protein levels of DRV-M-K195A-HA (I), ABLV-M-K205A-HA (J), DUVV-M-K195A-HA (K), and EBLV-M-K195A-HA (L) were analyzed by western blotting. Western blot data are representative of at least two independent experiments.

Then the K195 (or at the 205 site in ABLV-M) site of these lyssaviruses’ M proteins. The mutated M proteins were co-overexpressed with TRIM72 or empty vector in N2a cells for 48 h, and the levels of mutated M proteins were analyzed by western blot. As shown in Fig. 7I-7L, the levels of mutated M proteins were comparable between the TRIM72 over-expression group and the vector group, indicating that the mutation of the K195 (At the 205 site in ABLV-M) in lyssavirus M proteins impedes the degradation induced by TRIM72. Overall, the K195 (or at 205 site in ABLV-M) ubiquitination sites were partially conserved among lyssavirus M proteins and were essential for the degradation induced by TRIM72.

## DISCUSSION

The TRIM protein family is a large family consisting of more than 80 members, and TRIM proteins play important roles in innate and adaptive immune responses to defend against viral invasions [31]. Moreover, TRIM proteins can directly target viral proteins, leading to the degradation or functional inhibition of these viral proteins, thereby disturbing the viral lifecycle [31]. In this study, we have provided evidence demonstrating that TRIM72 restricts the release of lyssavirus by inducing the degradation of lyssavirus M protein in neuronal cells. First, we found that TRIM72 was up-regulated following lyssavirus infection in neuronal cells and mouse brains. Subsequently, we identified a direct interaction between TRIM72 and lyssavirus M, leading to the promotion of K48-linked ubiquitination of lyssavirus M at the K195 site, thereby facilitating lyssavirus M degradation via proteasome pathway, effectively restricting the release of lyssavirus (Fig. 8). Further study has revealed that the K195 site within M protein is partially conserved among lyssavirus M proteins. Importantly, these M proteins of lyssavirus that contain the conserved K195 site can be effectively degraded by TRIM72. This study is the first report demonstrating that lyssavirus M proteins undergo ubiquitination and degradation through the TRIM72-dependent proteasomal pathway.

**Fig. 8.**
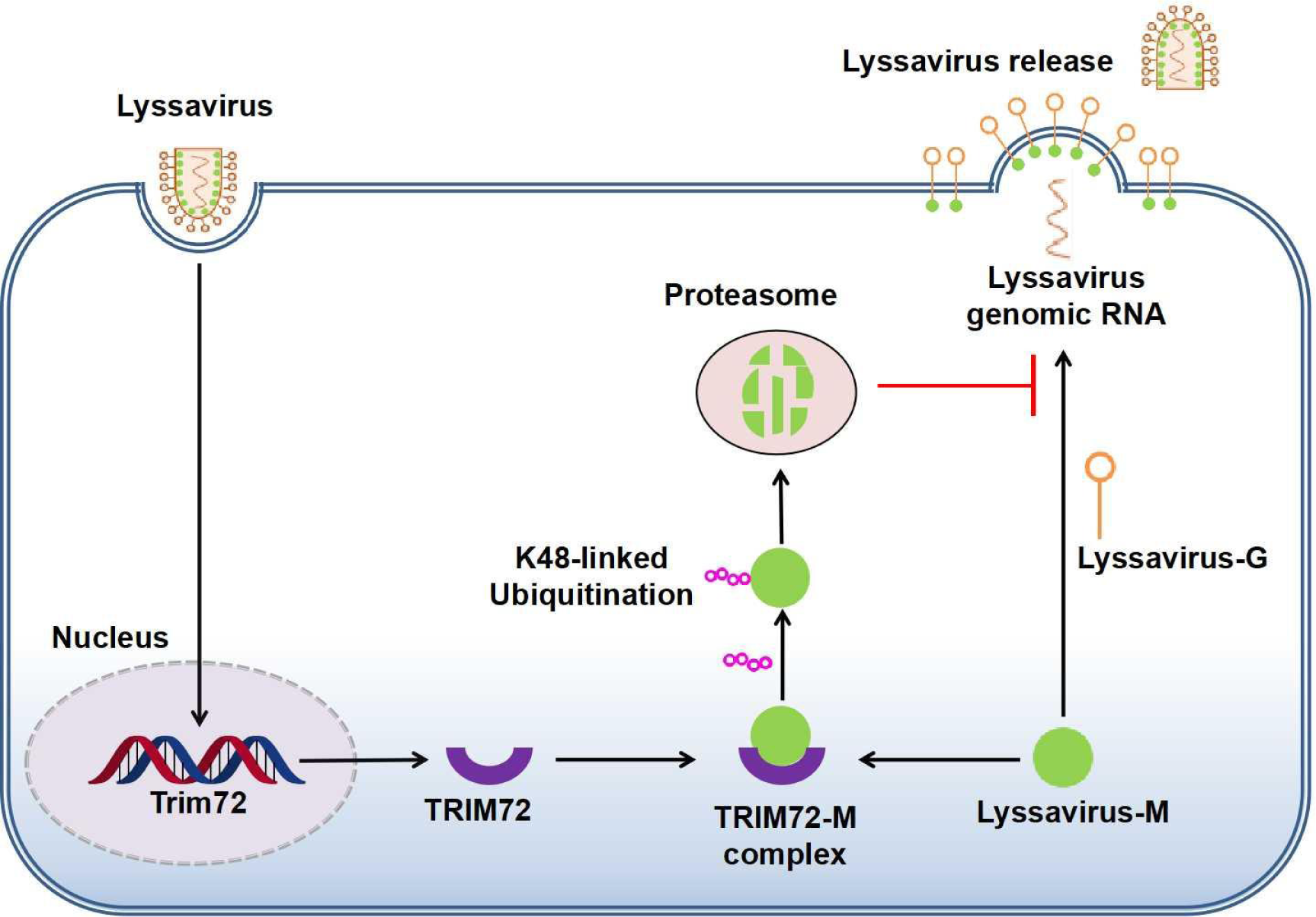
Model for TRIM72 restricts lyssavirus release by inducing K48-linked ubiquitination and proteasome degradation of the matrix protein.

TRIM72 plays an important role in membrane damage repair [32], and serves a protective role in many tissues, including neurons, heart, liver, kidney and muscle [33–38]. A previous study has indicated that TRIM72 attenuated LPS-induced neurotoxicity and neuroinflammation by inhibiting TLR4/NF-κB pathway both in vitro and in vivo [39]. A recent study has revealed that TRIM72 reduces muscle inflammation by promoting the inactivation of NLRP3 inflammasome [40]. Additionally, several studies have found a connection between TRIM72 and the influenza virus, demonstrating that TRIM72 protects mice from lethal influenza virus infection by suppressing inflammation [41, 42]. However. Until our study, there had been no reports of a direct interaction between TRIM72 and viral proteins. Our study here reveals a direct interaction between TRIM72 and lyssavirus M protein. Moreover, the interaction between TRIM72 and lyssavirus M promotes the degradation of lyssavirus M via a K48-linked ubiquitination modification in M at the K195 site. Recent studies have shown that TRIM72 is a potential E3-ligase enzyme [24, 25], and our findings in this study provide significant evidence to support this conclusion, confirming TRIM72 as an E3-ligase enzyme.

Ubiquitination is an important post-translational modification (PTM) that plays a key role in a wide range of cellular biological processes, including signal transduction [43], immune response [44], cancer [45], metabolism [46], etc. With a deeper understanding of ubiquitination, multiple forms of ubiquitination have been revealed, including K6-, K11-, K27-, K29-, K33-, K48- and K63-linked polyubiquitinations [47, 48]. Among the numerous polyubiquitination types, K48-linked polyubiquitination and K63-linked polyubiquitination have been the most extensively studied, multiple studies have proved that K48-linked polyubiquitination induces the degradation of target proteins and K63-linked polyubiquitination regulates the activation of target proteins [30, 47, 49]. K48-linked polyubiquitinations, which mediate proteasomal degradation, are the most abundant connection between ubiquitin proteins in cells [50]. The K48-linked polyubiquitination, responsible for the degradation of target proteins, plays essential roles in numerous biological processes, whether involving host or viral proteins. TRIM21, an interferon-inducible E3 ligase, has been shown to induce K48-linked ubiquitination and subsequent degradation of DDX41, thereby negatively regulating the innate immune response [51]. A most recent report has demonstrated that TRIM21 also restricts influenza A virus replication via a K48-linked degradation of the influenza A virus M1 protein [52]. These reports indicate that TRIM proteins could modulate the degradation of both host and viral proteins. TRIM72 has previously been reported to induce the K48-linked ubiquitination and degradation of FAK while regulating skeletal myogenesis [24]. However, there had been no prior reports of an association between TRIM72 and viral protein. In this study, we have demonstrated that TRIM72 directly interacts with lyssavirus-M and promotes K48-linked polyubiquitination at the lyssavirus-M K195 site, thereby promoting the degradation of lyssavirus M via the proteasome.

Lyssavirus M is the most abundant and smallest protein within the lyssavirus virion [53]. Lyssavirus M plays an essential role in the lyssavirus life cycle, with its primary functions encompassing viral assembly/budding and the regulation of the balance between transcription and replication [7, 8]. As a multiple-functional protein, lyssavirus M is involved in numerous biological processes during lyssavirus infection. M displays different functions via direct or indirect interaction with different host proteins and several important functions have been identified including defense host immune response, induce apoptosis and regulate lyssavirus replication [54]. NF-κB pathway is essential for defense against viral infection, and antiviral cytokines are produced once NK-kB is activated during virus infection. With this context, RelAp43, a member of the NK-kB pathway, has been found to participate in the innate immune response against lyssavirus infection [55]. Recent studies have revealed that lyssavirus M disturbs the NF-κB pathway by interacting with RelAp43 [9, 56]. Additionally, lyssavirus M was also found to interact with Jak1 and cooperate with lyssavirus P to modulate the Jak-Stat pathway, thereby restricting the expression of interferon-stimulated genes (ISGs) [57]. Lyssavirus induces apoptosis in neuronal cells during infection [58], and lyssavirus M contributes to this process via interacting with Cco1, ultimately decreasing the viability of Cco1, thereby disrupting the mitochondrial morphology and inducing apoptosis [59]. Furthermore, lyssavirus M facilitates its replication via interacting with host proteins, a recent study has shown that M interacts with ATP6V1A to promote lyssavirus uncoating [29].

TRIM proteins display an essential role in defending against virus invasion, while viruses also evolved functions to hijack the host ubiquitin system to facilitate their life cycle at different stages, including virus entry, virus replication, virus assembly and budding [60]. For example, TRIM7 induces K63-linked polyubiquitination on the E protein of Zika virus (ZIKV), the ubiquitination enhances ZIKV entry into cells via direct interaction with TIM1 a host cell receptor [61]. TRIM6 ubiquitinates the K309 site in the VP35 protein of Ebola virus (EBOV) and the ubiquitination facilitates the polymerase cofactor activity thereby enhancing the replication of EBOV [62]. Previous studies have shown that the ubiquitination of lyssavirus M which is regulated by the host is essential for lyssavirus budding [63, 64]. However, there are fewer reports on E3 ligases involved in lyssavirus M ubiquitination except for NEDD4 which is a membrane-localized ubiquitin ligase containing multiple WW domains that interact with the PPxY domain in the M protein of RABV [64]. Although the interaction sites between M and TRIM72-SPRY were conserved among lyssavirus M proteins (Fig. S4), it is worth noting that the degradation of lyssavirus M induced by TRIM72 is dependent on the ubiquitination at the K195 site, thus TRIM72 cannot induce the degradation of some lyssavirus M proteins due to the deficient of lysine at 195 site, such as LBV-M and MOKV-M. According to our results, there is a higher ubiquitination level in lyssavirus M following mutation at the M K195 site (Fig. 6C), indicating that there exist other ubiquitination sites in lyssavirus M. Although a previous study has identified a ubiquitination site at K60 of RABV M protein [65], additional ubiquitination sites may exist in lyssavirus M, necessitating further investigation.

In summary, our study has revealed a previously unknown role of TRIM72 during lyssavirus infection. We found that lyssavirus infection triggers an increase in TRIM72 levels in neuronal cells and mouse brains. Furthermore, we have identified a novel function of TRIM72, wherein it restricts lyssavirus release via directly interacting with lyssavirus M protein and facilitating the ubiquitination of lyssavirus’s M proteins at the K195 site, thereby promoting the degradation of lyssavirus M. The discovery in this study is significant because it sheds light on the direct interaction between TRIM72 and viral proteins, it also opens up the possibility that TRIM72 interacts and ubiquitinates other viral proteins from various viruses, which warrants further investigation. This research contributes to our understanding of how TRIM72 and other TRIM proteins play a role in defending against viral invasions and may inspire further research in this field.

## MATERIALS AND METHODS

### Cells

Cell lines N2a (mouse neuroblastoma cell, ATCC^®^ CCL-131), BSR (a clone of BHK-21, ATCC^®^ CCL-10) and SK-N-SH (human neuroblastoma cell, ATCC^®^ HTB-11) were obtained from American Type Culture Collection. Cells were cultured in a 37°C incubator with a humidified atmosphere containing 5% CO_2_. The growth media employed were DMEM (Bio-Channel Biological Technology Co., Ltd, BC-M-005-500ml) or RPMI1640, both of which were supplemented with 10% (vol/vol) fetal bovine serum (FBS, QmSuero/Tsingmu Biotechnology, Wuhan, China, mu001SR) and 1% penicillin-streptomycin (Bio-Channel, BC-CE-007-100ml).

### Primary neuron cells

Primary neuron cells were isolated from the cerebral cortex of 16-day C57BL/6 mouse embryos as described previously [66]. Briefly, the cerebral cortex was separated from the brains and then dissociated with 0.25% trypsin (wt/vol) at 37℃ for 30min. After centrifugation, the cells were resuspended in DMEM containing 5% FBS (Gibco) and subsequently passed through a 70 µm filter. After cell counting, the cells were seeded into 12-well plates (SORFA Life Science Co., Ltd, Beijing, China) that had been pretreated with poly-D-lysine and laminin (10 µg/ml) (Sigma, P7280). The cell supernatants were discarded at 5 h after seeding and replaced with Neurobasal plus Medium (Gibco, A3582901) containing 2% B-27 Plus (Gibco, A3582801).

### qPCR analysis

Total RNA was isolated from cells and tissues using TRIzol^®^ reagent (Aidlab Biotech, Beijing, China, 321728AX) following the manufacturer’s instruction. cDNAs were synthesized using HiScript^®^ II 1st Strand cDNA Synthesis Kit (Vazyme Biotech co., Ltd, R212-02), the genomic vRNA was synthesized using HiScript^®^ II Q RT SuperMix (Vazyme Biotech co.,ltd, R222-01) according to the manufacturer’s protocol. qPCR was performed using ChamQ Universal SYBR qPCR Master Mix (Vazyme Biotech Co., Ltd, Q711-02). The qPCR program consists of an initial denaturation step at 95℃ for 2 min for one cycle, followed by 40 cycles at 95℃ for 5 s and 60℃ for 30 s. The primer sequences used for qPCR were as follows (5’-3’):

Mouse Trim72-F GTTCTCACCGTGGTCATCGT
Mouse Trim72-R CAGCACCGCTACAGTCTTCT
hTrim72-F GACCCGCTGAGCATCTACTG
hTrim72-R AGCCACACTCTTCTCCTTGC
Mouse Gapdh-F CTACCCCCAATGTGTCCGTC
Mouse Gapdh-R TGAAGTCGCAGGAGACAACC
hGapdh-F AAGGTCATCCCTGAGCTAGAC
hGapdh-R GCAGGTTTTTCTAGACGGCAG
RABV N mRNA-F GATCGTGGAACACCATACCC
RABV N mRNA-R TTCATAAGCGGTGACGACTG
RABV-vRNA-F CTCCACAACGAGATGCTCAA
RABV-vRNA-R CATCCAACGGGAACAAGACT

### Western blot

Cells were lysed in lysis buffer (50 mM Tris pH=7.4, 150 mM NaCl, 0.25% Sodium deoxycholate, 1% NP40, 1 mM EDTA) containing 1x protease inhibitor cocktail (Biosharp, BL630B). After centrifugation at 12000 rpm/min for 10 min at 4 ℃, the supernatant was collected and SDS-PAGE loading buffer was added to the samples and boiled for 10 min. Total cell lysate samples were then separated on 8-12% SDS-PAGE gels and transferred to PVDF membranes (Millipore, ISEQ00010). Membranes were blocked with TBST containing 5% (w/v) non-fat dry milk at room temperature for 4 h or overnight in 4℃and probed with primary antibodies which were diluted with TBST and 5% (w/v) non-fat dry milk overnight in 4 ℃. The primary antibodies were against RABV N protein (prepared by our lab, 1:5000), RABV P protein (prepared by our lab, 1:5000), TRIM72 (Boster. A06982-2), Flag tag (MBL, M185-3L, 1:10000), HA tag (MBL, M180-3, 1:10000), Myc tag (MBL, M192-3, 1:10000), or GAPDH (ProteinTech, 60004-1-Ig, 1:5000). Membranes were probed with HRP-conjugated goat anti-mouse secondary antibodies (Boster, Wuan, China, BA1051), goat anti-rabbit secondary antibodies (Boster, BA1055, 1:6000), goat anti-mouse IgG light chain secondary antibodies (Abbkine, A25012, 1:5000), goat anti-mouse IgG Heavy chain secondary antibodies (Abbkine, A25012, 1:5000), then developed using BeyoECL Star kit (Beyotime, P0018A). Images were captured with an Amersham Imager 600 (GE Healthcare) imaging system.

### Trim72 specific siRNA

Trim72-specific siRNAs were designed and synthesized by Ribobio Co., Ltd The siEDAL sequences are listed below:

siTrim72-1: 5’-GCACGCCUCAAGACACAGC-3’;
siTrim72-2: 5’-UCAAAUUCCAGGUGUGGAA-3’;
siTrim72-3: 5’-GCUUUCUAUGAUGCGAGCA-3’.

### Rescue of rRABVs

The recombinant rRABV-Trim72, rRABV-Trim72(−) and mutant rRABV-M-K195A were cloned from RABV strain challenge virus standard-B2c (CVS-B2c) and constructed as described previously [67, 68]. Briefly, BSR cells were seeded in 6-well plates (SORFA Life Science Co., Ltd, Beijing, China) and transfected with 2 µg of a fully infectious clone, 0.5 µg of pcDNA-N, 0.25 µg of pcDNA-P, 0.15 µg of pcDNA-G and 0.1 µg of pcDNA-L by using Jetprime polyplus transfection reagent (Polyplus, 114-15) according to the manufacturer’s instructions. Four days post-transfection, supernatants were harvested and examined for the presence of rescued viruses using FITC-conjugated anti-RABV P antibodies (Prepared by our lab).

### Virus challenge experiment in mice

6-week-old female C57 BL/6 mice were infected intracerebrally with 200 fluorescence focus units (FFU) RABV (CVS-B2c) or mock infected, and the mouse brains were collected at 6 d post-infection for qPCR and western blot analysis.

8-week-old female C57 BL/6 mice were randomly divided into four groups (n=10). 100 FFU of recombinant RABV (rRABV), rRABV-Trim72 and rRABV-Trim72(−) were injected into mice via intranasal individually, and the same volume of DMEM was injected into control mice. Survival percent, body weight changes and clinical score were recorded daily for 21 days.

### IHC staining

Slices of mouse brains were prepared as described in our previous study [68]. Briefly, mice were intracardially perfused with PBS, and the brains were then extracted and fixed in 4% paraformaldehyde at 4℃ for 16 h. Subsequently, the brains were embedded in paraffin and sectioned into slices. After dewaxing and dehydration, the slices were stained with a RABV-P polyclonal antibody (Prepared by our lab) overnight at 4℃. Following incubation with HRP-conjugated anti-rabbit secondary antibodies (Servicebio, GB23303, 1:200) were incubated, the slices were treated with diaminobenzidine (Servicebio, G1211) for color development. Finally, an XSP-C204 microscope (CIC) was used for photography and analysis.

### VLP analysis

VLP system was reconstructed as previously described. Briefly, empty vector or pCA-TRIM72 together with pCA-M and pCA-G (ratio 6:1) were co-transfected into HEK 293T cells for 48 h. Cells were collected for the analysis of the protein expression level by western blotting, and the supernatants were collected and centrifugated at 4 ℃ for 30 min (5200 rpm/min). Then the supernatants were collected again and centrifugated at 4℃ for 30 min (30000 rpm/min). The pellets were then collected and used for the analysis of VLP production by western blotting.

### Co-immunoprecipitation (Co-IP)

The pretreated cells were washed 3 times with PBS and then lysed in lysis buffer (50 mM Tris pH=7.4, 150 mM NaCl, 0.25% Sodium deoxycholate, 1% NP40, 1 mM EDTA) containing 1x protease inhibitor cocktail (Biosharp, BL630B). After centrifuging at 12000 rpm/min for 10 min at 4 ℃, the supernatant was collected. Anti-flag mAb-magnetic agarose (MBL, M185-10) or Anti HA-tag mAb-magnetic agarose (MBL, M180-10) was then added to the samples and incubated for 4 h at 4 ℃. After incubation, the samples were washed 3 times with lysis buffer (5 min/time), and then the magnetic agarose was collected and resuspended in PBS. Then SDS-PAGE loading buffer was added to the samples and boiled for 10 min. The supernatants were collected and used for western blot analysis.

## Author contributions

Conceived and designed the experiments: L.Z., B.K.S. and M.Z.

Performed the experiments: B.K.S. and J.X.Z.

Analyzed the data: B.K.S., J.X.Z., Z.F.F., L.Z. and M.Z.

Wrote the paper: B.K.S., L.Z. and M.Z.

## Acknowledgements

This study was supported by the National Natural Science Foundation of China (grant number 32102648 to B.K.S.), the Fundamental Research Funds for the Central Universities (grant number 2662023PY005 to L.Z.), the National Key Research and Development Program of China (grant number 2022YFD1800100 to M.Z.), the China Postdoctoral Science Foundation (grant number BX2021109 to B.K.S., grant number 2021M691170 to B.K.S.).

## Ethical approval

Female C57 BL/6 mice (8-week-old) were purchased from the Hubei Center for Disease Control and Prevention, Hubei, China, and were housed in the animal facility at Huazhong Agricultural University in accordance with the recommendations in the Guide for the Care and Use of Laboratory Animals of Hubei Province, China. All experimental procedures involving animals were reviewed and approved by The Scientific Ethic Committee of Huazhong Agricultural University (permit number HZAUMO-2019-086).

## Statistical analysis

Statistical analysis was performed using GraphPad Prism 6. The significance of differences was analyzed with Student’s t-test or Two-way ANOVA test. Survival percent was analyzed by log-rank (Mantel-Cox) test. *P <0.05, **P <0.01, ***P < 0.001, ****P < 0.0001.

## Supplementary figures

**Fig. S1.**
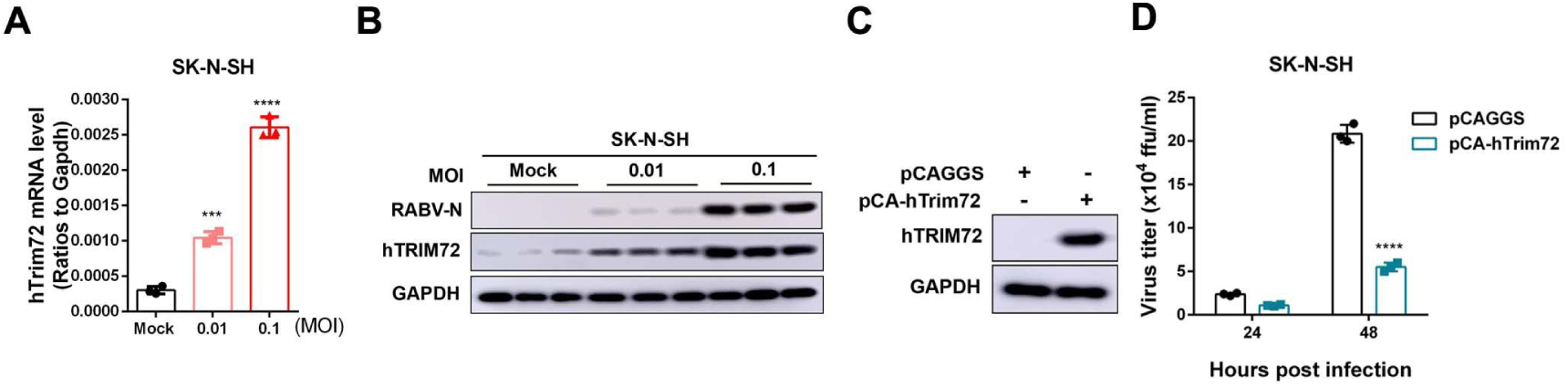
RABV infection induces hTRIM72 up-regulation in SK-N-SH cells and hTRIM72 reduced RABV replication in SK-N-SH cells. (A-B) SK-N-SH cells were infected with RABV at different MOI for 36 h. The mRNA level of TRIM72 was analyzed by qPCR (G), and the protein levels of TRIM72 and RABV-N were analyzed by western blotting (H). (C) Empty vector (pCAGGS) or hTRIM72 over-expression vector (pCA-hTrim72) were transfected into SK-N-SH cells respectively for 48 h, then hTRIM72 level was analyzed by western blotting. (D) Empty vector or hTRIM72 over-expression vectors were transfected into SK-N-SH cells respectively for 12 h, then infected with RABV (MOI=0.01) for the indicated time, and the viral titers in the supernatant were analyzed. Statistical analysis of grouped comparisons was carried out by student’s t-test (*P < 0.05; **P<0.01; ***P<0.001; ****P<0.0001). The bar graph represents means ± SD, n = 3. Western blot data are representative of at least two independent experiments.

**Fig. S2.**
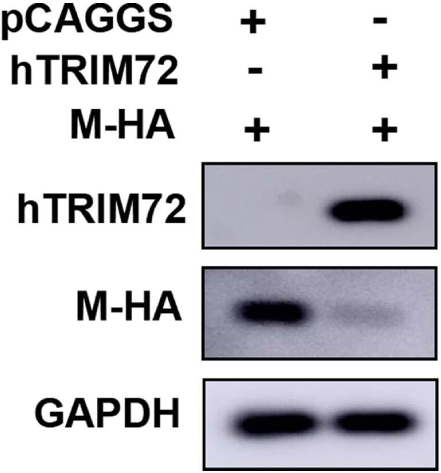
hTRIM72 induces the degradation of M. pCAGGS or pCA-hTrim72 together with pCA-M-HA were co-transfected into SK-N-SH cells respectively for 48 h, and the protein levels of M and hTRIM72 were analyzed by western blotting. Western blot data are representative of at least two independent experiments.

**Fig. S3.**
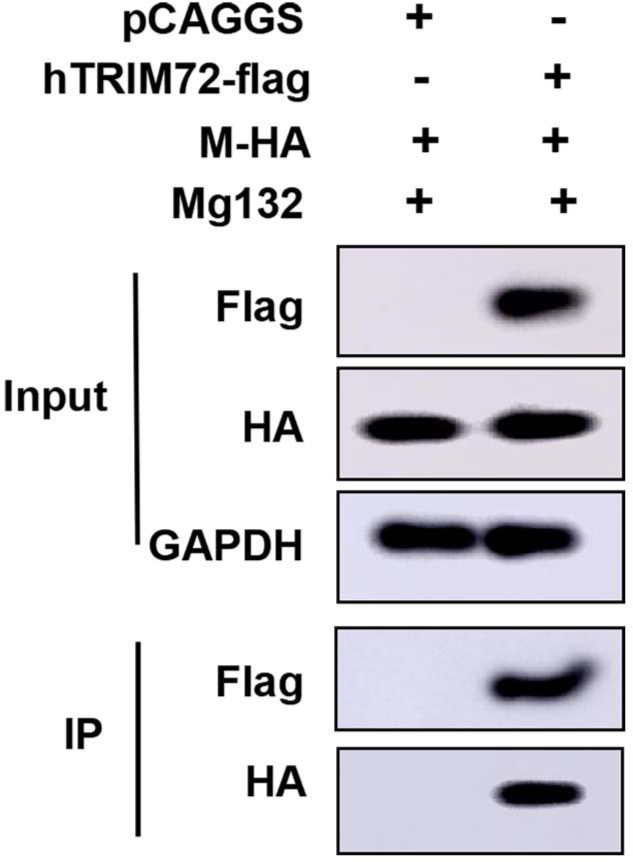
hTRIM72 interacts with RABV-M. pCAGGS or pCA-hTrim72-flag together with pCA-M-HA were co-transfected into N2a cells respectively. Then Mg132 (10 µM) was treated, and Co-IP assays were performed with anti-flag antibody post-transfection for 48 h. The protein levels of TRIM72-flag and M-HA were analyzed by western blotting. Western blot data are representative of at least two independent experiments.

**Fig. S4.**
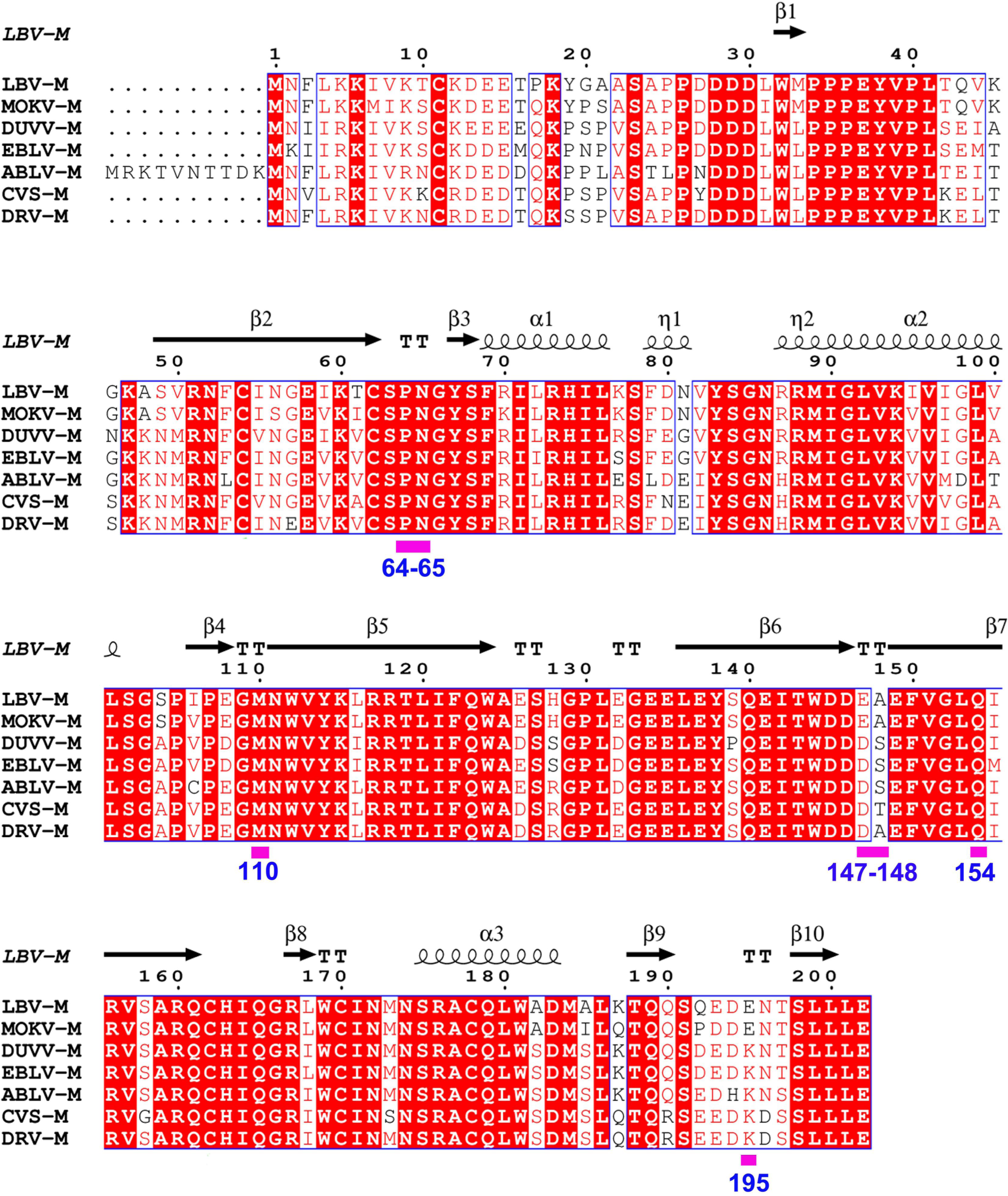
Lyssavirus M proteins’ sequences were compared. The protein sequences of lyssavirus M were compared and analyzed with ESPript 3.0 online software (https://espript.ibcp.fr/ESPript/cgi-bin/ESPript.cgi).

**Fig. S5.**
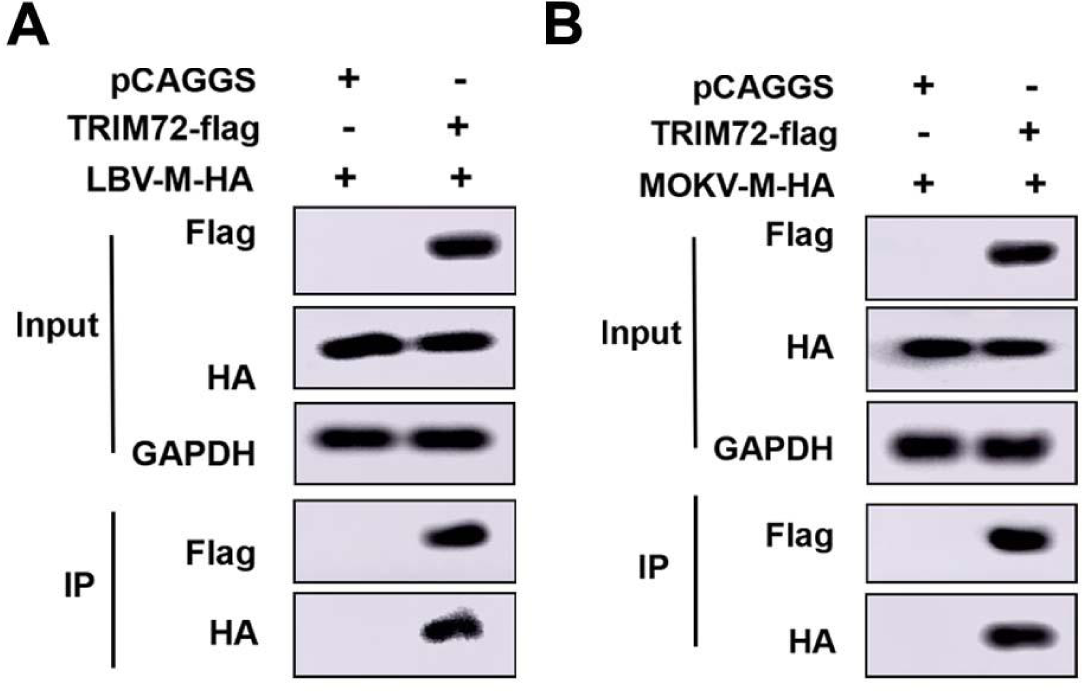
TRIM72 interacts with LBV-M and MOKV-M. (A-B) HA-tagged lyssavirus M proteins (LBV-M-HA and MOKV-MHA) together with empty vector or TRIM72-flag were co-overexpressed in N2a cells respectively. Co-IP assays were performed with anti-flag antibody post-transfection for 48 h and protein levels of LBV-M-HA (A), MOKV-M-HA (B), were analyzed by western blotting. Western blot data are representative of at least two independent experiments.

